# Structure of Macrolide Efflux Protein (MefA/E) in *Streptococcus pneumoniae*: An in silico approach

**DOI:** 10.1101/2020.07.21.213744

**Authors:** Sreeram Chandra Murthy Peela, Jyoti Sharma, Sujatha Sistla

## Abstract

**Background:** Macrolides are one of the commonest antibiotics used to treat bacterial respiratory tract infections. Resistance to this class of antibiotics is on the rise and is mediated by macrolide efflux (MefA/E) protein as one of the mechanisms. Despite its importance, the structure of this protein is not known yet.

**Methods:** The publicly available MefA/E protein sequence was used to model the structure. Modelling was performed in I-TASSER, and the model was further refined. Its orientation in a membrane was studied using OPM server.

**Results and conclusions:** The structure of MefA/E resembled that of Major Facilitator Superfamily (MFS) proteins, with 13 transmembrane helices. It had a V-shaped conformation, with the wider part towards the outer membrane layer.

## Introduction

Macrolides are one of the commonly prescribed antibioitcs for treating respiratory infections. They act by inhibiting bacterial protein synthesis and are bacteriostatic or bactericidal depending on their concentration and the bacteria. While it continues to be given as emperical therapy, there is a substantial rise of resistance to this group of antibiotics globally in pathogens like *Streptococcus pneumoniae* that cause lower respiratory tract infections. In fact we estimate the resistance to macrolides in S. pneumoniae to be 35.1%, and two mechanisms contributed to the resistance - efflux mediated (encoded by MefA/E) and ribosomal methylation (mediated by ermB) (1). Also there are less common mechanisms like mutations in 23S-rRNA regions like L4 and domain V (2). In our study, efflux mediated macrolide resistance was detected in 34% of the 47 isolates resistant to macrolides.

The macrolide efflux protein of pneumococcus belongs to the Major Facilitator Superfamily (MFS) and is 403 amino acids long. The protein shares >80% similarity with macrolide efflux protein MefA of *S. pyogenes* and sometimes both these proteins are represented as MefA/E (3). Similar homologues were detected in other bacteria as well (MefC in *Vibrio*, MefI in *S. pseudopneumoniae*, MefO in *S. dysgalactiae*, etc). The gene is usually present on Mega element, Tn2009, and Tn2010 transposons (4). These, in turn, carry an ABC-type transporter protein called Mel or MsrD which has two fused nucleotide-binding domains but no transmembrane domains. The genes MefE and Mel are usually transcribed together and have macrolides as their inducers, especially the 14- and 15-membered macrolides (5,6). The co-expression of both these genes is required for macrolide efflux in pneumococci, and they interact synergistically in *E.coli* (a physical interaction between MefE and Mel is detected in *E.coli*). The expression of these genes is regulated by transcription attenuation (7). The anti-attenuation of transcription in the presence of macrolides leads to the expression of these genes.

Despite its significance, the structure and mechanism of MefA/E is not studied in detail. The present study was thus aimed to model the structure of MefA/E using computational methods.

## Materials and Methods

The MefA/E protein sequence with GenBank accession number WP_000417519.1 was selected for the modeling. The initial search in the PDB database was performed to check for homologous sequences. When no similar sequences were found in the PDB database, and as the protein is known to be membrane-associated, homology modeling and threading was performed to predict the 3D structure of this protein. The structure prediction was performed in the I-TASSER server using default parameters (8). Once the models were generated, their validation was performed using a Ramachandran plot, ProQM, and QMEANBrane programs of SWISSPDB server (9–11). Best models were selected and further validated using ERRAT plot in SAVES web server (12). Any model refinement was performed in ModLoop and GalaxyRefine servers and new models were revalidated (13,14). The process is iterated until the best model was obtained. Finally, 2000 steps each of the steepest descent and conjugate gradient were performed in Swiss PDB viewer (SPDBV) using GROMOS96 forcefield and in vacuo. The orientation of the protein in the membrane was estimated using OPM (Orientation of Proteins in Membranes) server (15). All the final images were downloaded from respective sites and protein structure visualisation was performed in DISCOVERY STUDIO.

## Results

The predicted structure of the MefE protein predominantly consisted of α – helices with a central cavity (figure 1). The predicted model had a TM-score of 0.70 ± 0.12 and a cluster density of 0.2594 using 1992 decoys. The accuracy of the predicted model was acceptable when analysed using normalised B-factor and residue-specific error in I-TASSER suite (figure 2). The initial structure had a few outliers and errors when assessed using Ramachandran and ERRAT plots respectively. Subsequent refinement of protein structure removed these errors and the overall quality factor of the final protein was 99.496 with no outliers (figure 3). The global S-score in the ProQM server was 0.605 while the local S-scores ranged from 0 to 0.75. The predicted structure had membrane insertion energy within the range of transmembrane proteins with most of the residues having local quality > 0.5 (figure 4). The structure, when embedded in the membrane, had a depth/hydrophobic thickness of 30.6 ± 1.1 Å and a tilt of 8.0± 2.0° with ΔG_transfer_ of −90.4 kcal/mol (figure 5). The embedded and transmembrane residues are shown in the table where the total number of the transmembrane segments were 13 (table 1).

**Table 1:** Predicted embedded and transmembrane residues in MefE modeled structure

| Subunits | Tilt | Segments |
| --- | --- | --- |
| A | 22 | Embedded residues:<br>12-36,39,43-64,67,76-97,100-124,140-162,164-187,216,224-246,250,253-274,288-329,331-332,348-392 |
| A | 7 | Transmembrane segments:<br>1(12- 36), 2(43- 64), 3(77- 97), 4(102- 124), 5(140- 162), 6(167- 186), 7(224- 233), 8(237- 247), 9(254- 274), 10(290- 305), 11(310- 329), 12(348- 373), 13(374- 391) |

**Figure 1:**
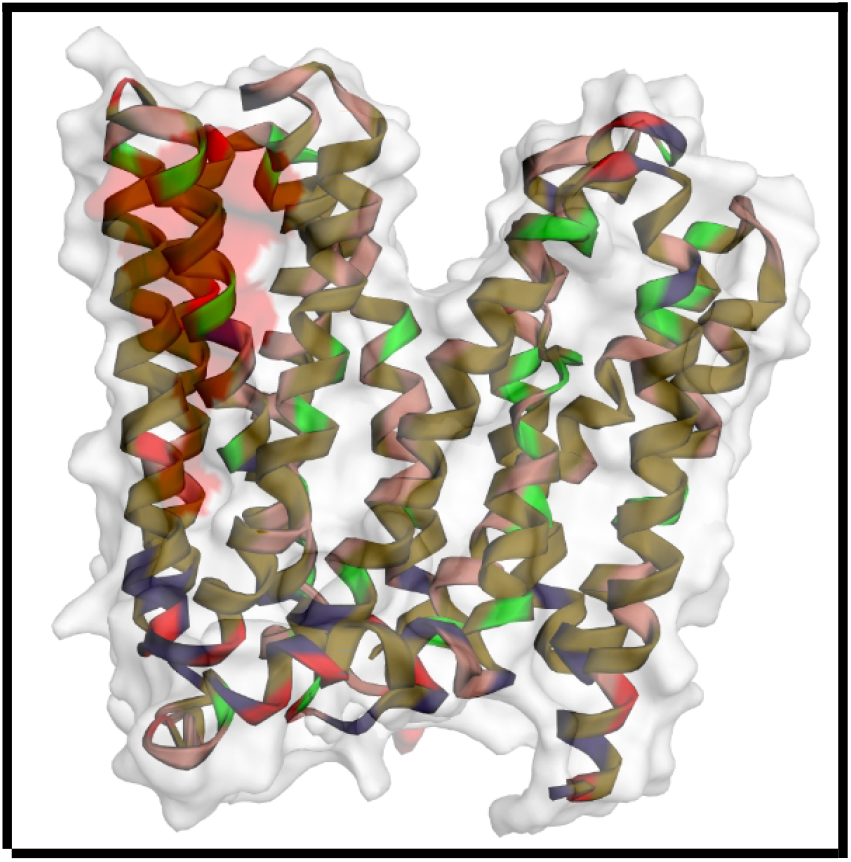
Modeled MefA/E protein Structure

**Figure 2:**
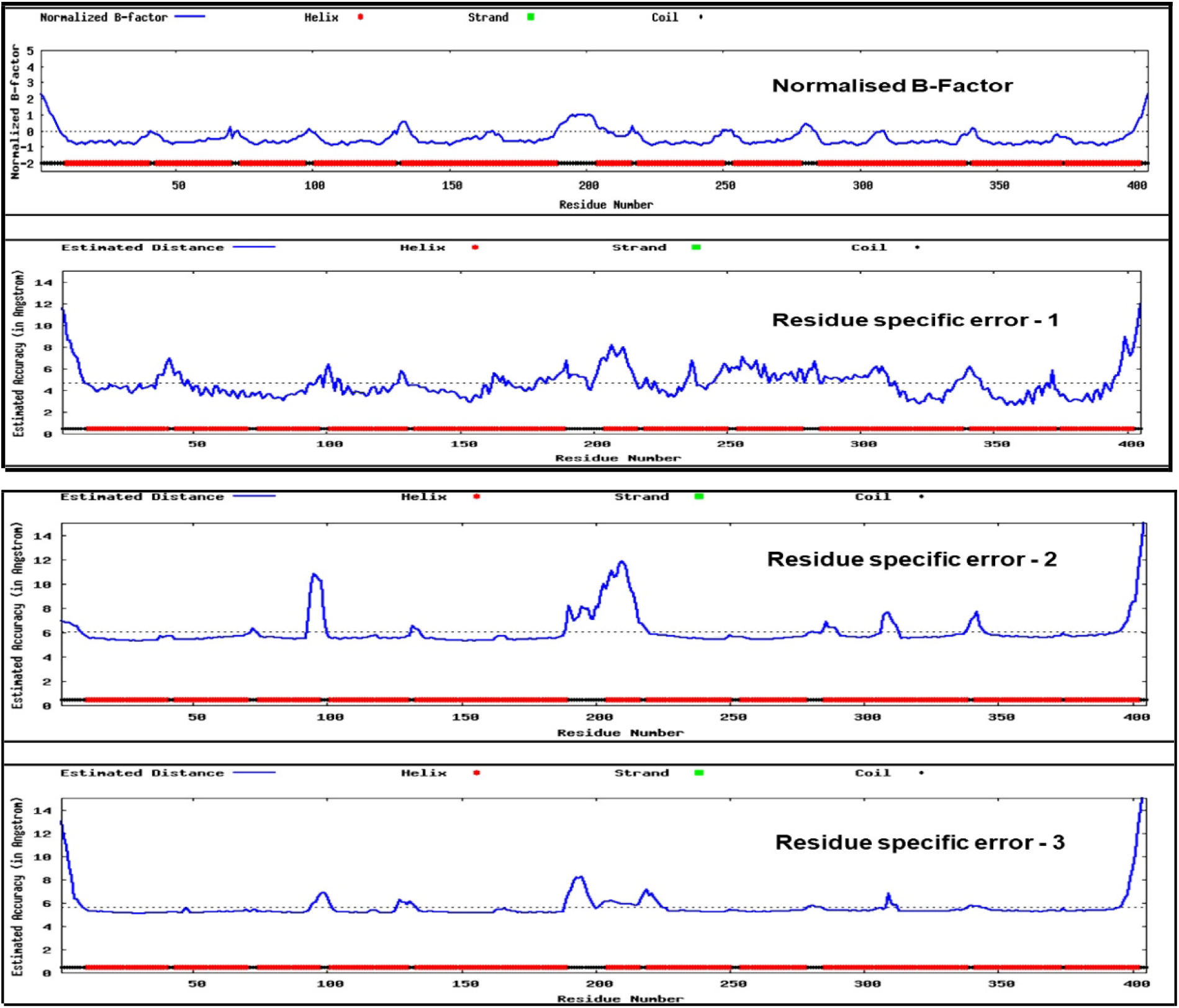
Model evaluation by I-TASSER suite by calculating the normalised B-factor and residue-specific error.

**Figure 3:**
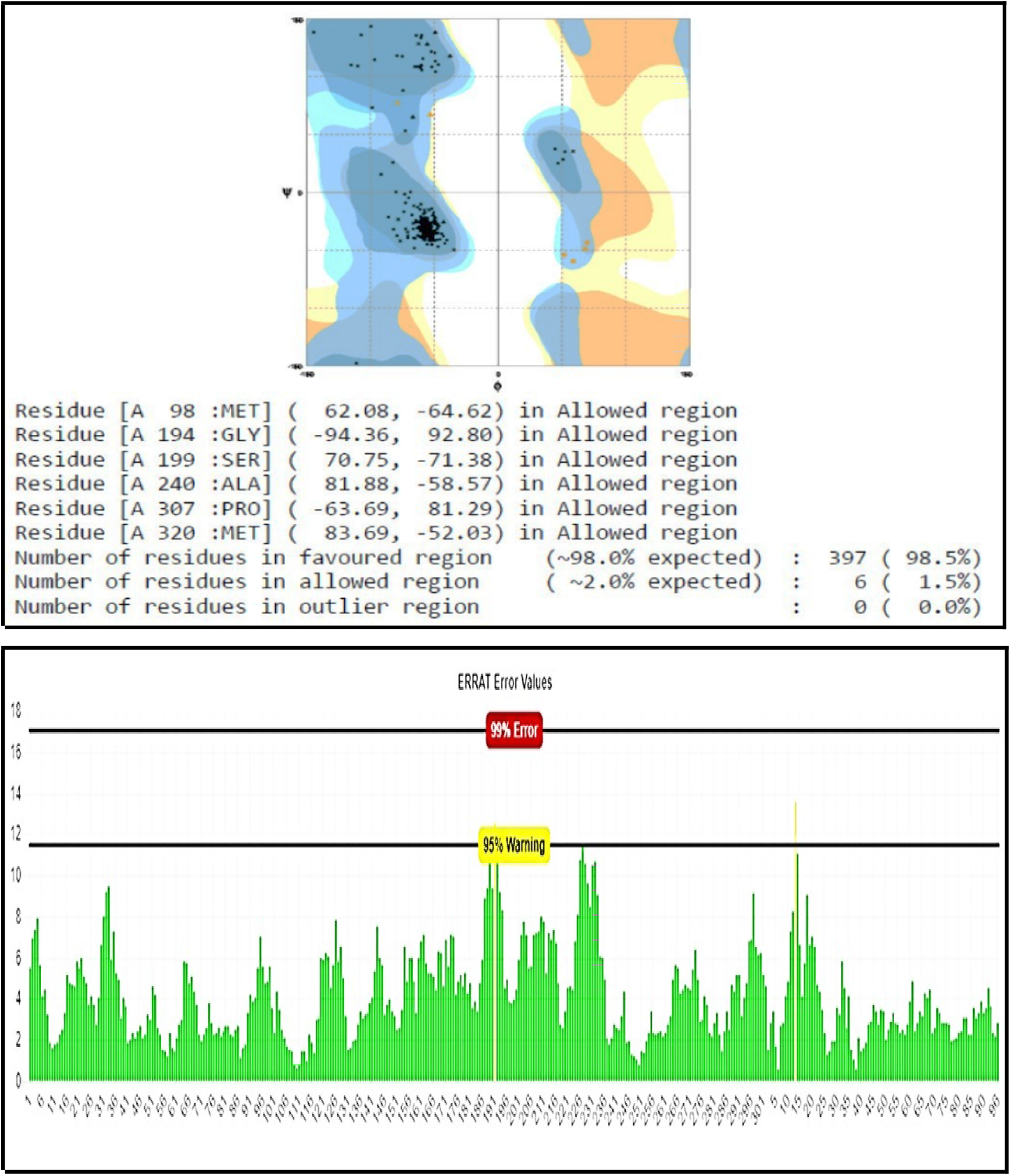
Ramachandran plot and ERRAT plot for validation of predicted structure.

**Figure 4:**
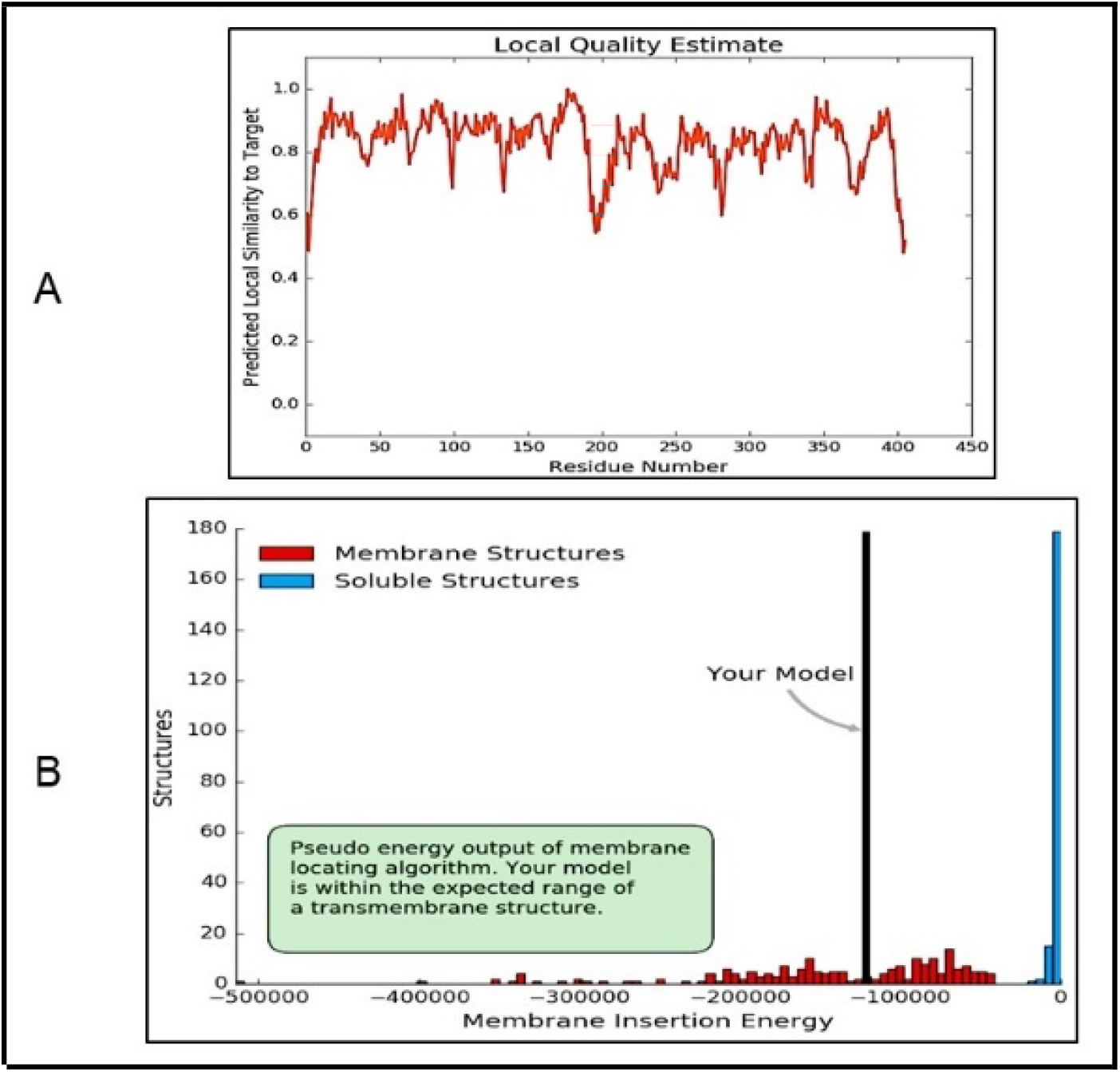
QMEANBrane assessment of quality estimate and model insertion energy.

**Figure 5:**
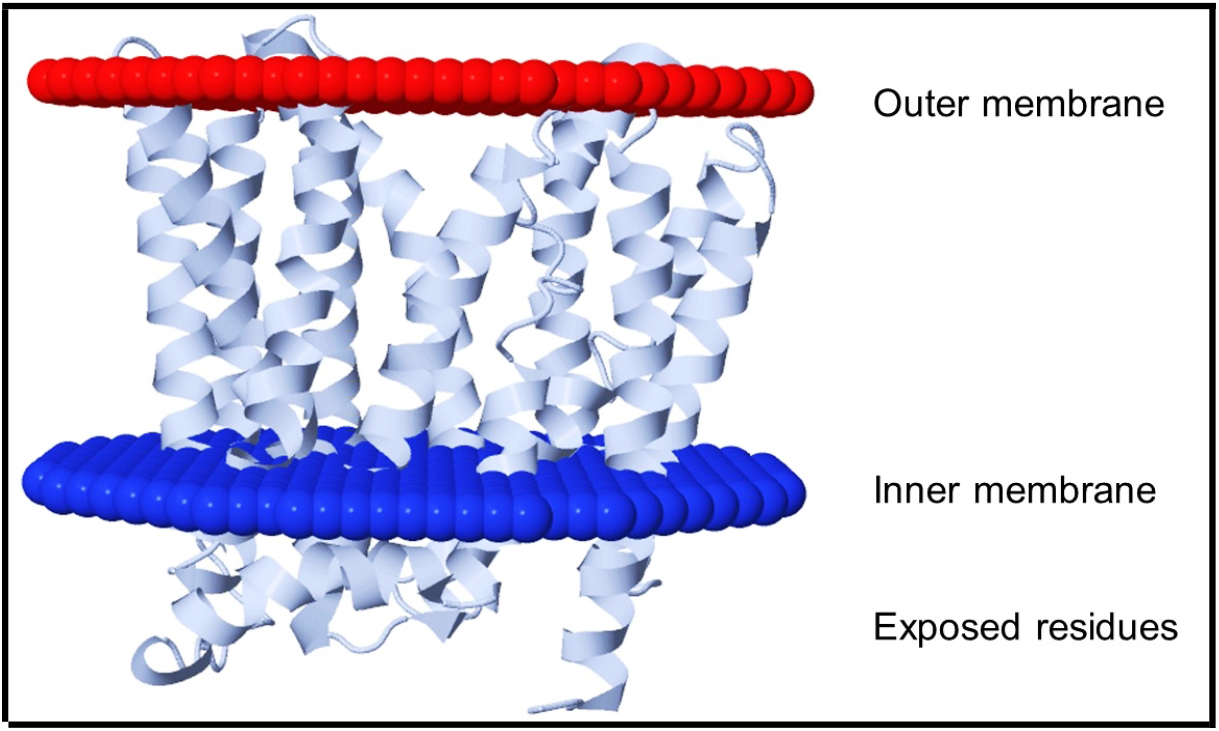
Orientation of *MefA/E* in the cell membrane. The opening of the channel is wider towards the outer membrane.

## Discussion

The macrolides are one of the most commonly prescribed drugs to treat upper respiratory infections. They are even administered in cases of pneumonia caused by the pneumococcus and are the group A drugs as per CLSI 2019 (needs to be reported for all the isolates). Among the macrolides, azithromycin is commonly available as an over the counter antibiotic. It has a once-daily oral mode of administration and the therapy usually lasts for three days. Due to its abuse, the spread of macrolide resistance has been rapid in recent years. In many centers across the world, the macrolide resistance has been increasing and currently is in the range of 20% to 35% in India (1). The spread of macrolide resistance is mediated commonly by two mechanisms – ribosomal methylation (*ermB*-mediated) and efflux of the drug (MefA/E-mediated) (7). These genetic mechanisms are not limited to pneumococcus but are also common among the streptococci like *S. pyogenes*. The mechanism of action of these two different methods was not studied at the molecular level.

To study the molecular mechanism of MefA/E, no crystal structures were available as of now as per an initial search in PDB using NCBI-BLASTP tool. Hence a combination of homology modeling and threading was employed to predict the structure of MefA/E. For this purpose, a multispecies protein sequence with GenBank accession number WP_000417519.1 was selected for modelling. The models were generated using I-TASSER web server (8). The I-TASSER uses C-score to predict the global (overall) structure accuracy and ResQ for residue-specific local quality. Typically the C-score ranges from −5 and 2 with a higher score predicting more accurate models. In the present study, the model selected had a C-score of −0.14. The TM-score and RMSD are correlated with the C-score and are the standards to predict the accuracy of modeled structures when original structures are available. In cases where original structures are not available, these values are predicted from the C-score. While the RMSD describes the global structure differences, the TM-score measures the local structural variations. If the TM-score is >0.5, the topology of the modeled structure is more accurate. The TM-score of the predicted MefA/E protein was 0.7±0.12 which means that the topology is highly accurate.

The modeled structure was further assessed using a Ramachandran plot, ERRAT, and QMEANBrane webservers. The initial model had 10 residues in the outlier region with an overall quality factor of 97.23 and high local quality scores. To remove the outliers, the model was refined used the GalaxyRefine webserver. The refined structure had no outliers in the Ramachandran plot and an overall quality factor of 99.496. This model was further validated using ProQM in which the global and local scores were 0.605 and 0-0.75 respectively. While the Ramachandran plot and ERRAT work well for globular proteins, the validation of a membrane protein by these methods is inaccurate. The ProQM is one of the membrane protein quality assessment tool, and displays the quality score called S-score and ranges from 0 and 1 (worst and best models respectively). As seen from above, the predicted MefA/E model had S-score of 0.605 indicating a good model. The final model had 13 transmembrane helices similar to those found in MFS family of proteins (12 transmembrane helices). A central cavity is present throughout the protein and maybe the channel for drug efflux. The 13 helices are embedded in the lipid bilayer as evidenced by the OPM server. The general structure of MFS family consists of two domains with a central pore. The structure of MefA/E predicted follows the same pattern of having a central cavity surrounded by helical chains.

These proteins operate via a rocket switch mechanism with two intermediate states – the inward-facing (Ci) and the occluded (Co) conformations. The MefA/E is similar to the Ci conformation of GlpT with two domains (16). While the structure has been successfully modeled, the structure of Mel which is known to associate with MefA/E and interactions with the drug molecules has to be performed.

## Conclusion

The macrolide efflux protein (MefA/E) was successfully modeled in silico and validated using a variety of computational tools. Its orientation in the membrane was found to be similar to that of MFS super-family proteins in having 13 transmembrane rings and having a V-shaped conformation.

